# DMC1 and RAD51 bind FxxA and FxPP motifs of BRCA2 via two separate interfaces

**DOI:** 10.1101/2023.10.05.561053

**Authors:** Simona Miron, Pierre Legrand, Sari E. van Rossum-Fikkert, Dejan Ristic, Atifa Badar, Roland Kanaar, Sophie Zinn-Justin, Alex N. Zelensky

## Abstract

In vertebrates, the BRCA2 protein is essential for meiotic and somatic homologous recombination (HR) due to its interaction with RAD51 and DMC1 strand exchange proteins (recombinases). The interaction is mediated by FxxA and FxPP motifs, whose defining feature is the invariant phenylalanine. The FxxA motifs, present in the eight BRC repeats in the central region of BRCA2, compete with the FxxA motif of the linker region of RAD51 that is responsible for recombinase self-oligomerization. In vitro, BRCs disrupt RAD51 nucleoprotein filaments, but they are essential for RAD51 function in the context of the full-length BRCA2 protein. The role of the FxPP motifs is poorly studied but they also contribute to BRCA2 function in cells. In particular, the C-terminal TR2/CTRB domain of BRCA2, which contains an FxPP motif, is required for stabilization of RAD51 filament and replication fork protection. We recently found that deletion of the BRCA2 PhePP domain, which contains another FxPP motif, disrupts DMC1 but not RAD51 function in meiosis. Here we provide a mechanistic explanation for this phenotype by solving the crystal structure of the complex between DMC1 and the PhePP domain of BRCA2. Our structure reveals that, despite sequence similarity, the A-motifs (FxxA) and P-motifs (FxPP) bind to distinct and contiguous sites on the recombinases. The PhePP P-motif binding site is mostly located at the ATPase domain surface of a DMC1 monomer, but also extends to the linker region of the adjacent monomer, thus engaging two adjacent protomers in the DMC1 oligomer. Our structural analysis provides a mechanism explaining how PhePP favors the formation of the DMC1 nucleoprotein filament and stabilizes it. It corroborates and explains the stabilizing effect of the P-motif from BRCA2 TR2/CTRB on RAD51.

## Introduction

BRCA2 is essential for homologous recombination (HR) in somatic cells ^1,2^ and during meiosis ^3–9^. This function is attributed to its interaction with the recombinases: the ubiquitous RAD51 ^10,11^ and the meiosis-specific DMC1 ^12–16^. BRCA2 facilitates the loading of RAD51 and DMC1 on single-strand DNA (ssDNA) produced by end resection during HR. Thus, it controls the formation of helical nucleoprotein filaments ^17^, which catalyze homology search and strand exchange — the key events in HR. BRCA2 acts as an HR mediator ^18^, helping to evict the competing ssDNA-binding proteins and to direct filament formation to the correct substrate. Less detail is known about BRCA2 in meiosis, e.g., whether BRCA2 coordinates both recombinases directly, or affects DMC1 indirectly via its effect on RAD51 ^13,19^. We engineered a mouse with an internal deletion of *Brca2* exons 12-14, the region we called “the meiotic domain of BRCA2” because only meiotic functions were abolished ^20^. In the females, we found that DMC1 foci became undetectable, but RAD51 foci remained. Thus, BRCA2 indeed directly controls DMC1 and has a dedicated domain for it, lost in our mouse strain. The deleted region contains the so-called “PhePP motif”, which was previously shown to bind DMC1 ^14^, but whose relevance was not clear ^21,22^. We confirmed that PhePP binds to DMC1 and not to RAD51 ^14,22^, but the details of the interaction and why it is required for DMC1 function remained unresolved. In this study we aimed to address this using structural and biochemical methods.

The human BRCA2 PhePP motif, corresponding to the sequence FVPP, was discovered in vitro ^14^, but not studied further, except in a mouse model that questioned its relevance for mammalian meiotic HR ^21^. It is similar to the much better characterized phenylalanine-containing FxxA motif of the BRC repeats, which represents a conserved feature of BRCA2 in eukaryotes and binds RAD51 and DMC1 (Figure 1A). Other motifs with a FxxP sequence were detected in BRCA2 from vertebrates, invertebrates, plants and fungi ^14^. In Brh2, an ortholog of BRCA2 from fungus *Ustilago maydis* and a crucial model ^23–25^, such motifs act synergistically with BRCs, leading to suggestion of similarity and redundancy between the motifs FxxA and FxxP ^26,27^. The FxxP motifs interact with RAD51 (DMC1 is absent from *Ustilago maydis*) thus contributing to DNA repair proficiency ^26^. In vertebrates, a motif called TR2 ^28^, CTRD ^29^ or CTRB ^30^ that binds to RAD51 was detected in the C-terminal disordered region of BRCA2. It has a stabilizing effect on RAD51-DNA complexes, counteracting BRCs ^31,32^. Research on this C-terminal site was focused on S3291, which can be phosphorylated by CDKs at the G2-M transition ^28,30–35^.

**Figure 1.**
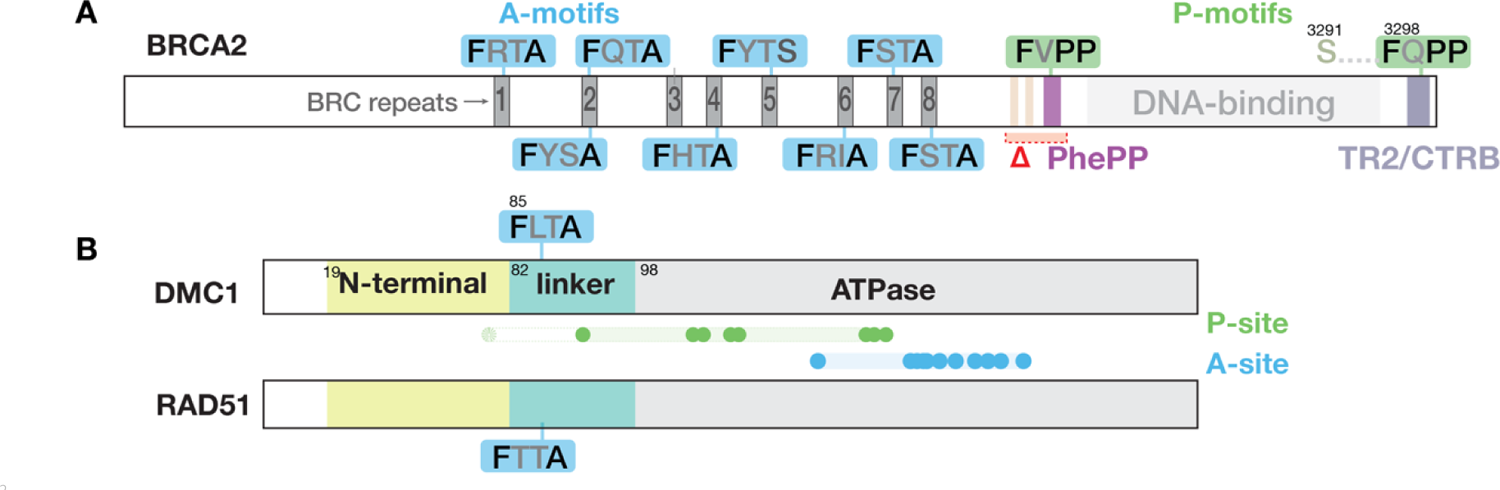
Definition of recombinase-binding motifs. The figure depicts schematically the motifs and domains of (A) BRCA2, and (B) the recombinases DMC1 and RAD51. We refer to the conserved FxPP (green) and FxxA (blue) motifs as P- and A-motifs, and regions on the recombinases binding to P- and A-motifs (dots: residues defining the sites) as P- and A-sites, as detailed in the results section. Residue numbers indicated for the domain boundaries and key amino acid positions relate to human proteins. Region deleted in the *Brca2^Δ^*^12–14^ mouse is shown in red (Δ) in panel A.

Phosphorylation abolishes the stabilizing effect, suggesting an HR control mechanism. A recent study revealed that rather than S3291, the adjacent highly conserved F3298 is essential for the CTRD-RAD51 interaction, and the surrounding conserved motif matches the FxPP consensus of the PhePP motif. Thus, whereas our *Brca2^Δ^*^12–14^ mouse model supports a meiotic function for the interaction between the BRCA2 FxPP motif encoded by exon 14 and DMC1, another BRCA2 FxPP motif encoded by exon 27 binds to RAD51, suggesting that interactions between FxPP motifs and recombinases share structural and functional properties ^22^.

The determinants of the FxxA interaction with RAD51 were studied in a fusion protein hBRC4∼RAD51-ATPase domain ^36^, to overcome the tendency of purified human RAD51 to aggregate during crystallization attempts. The crystal structure of this protein revealed that the invariant phenylalanine from the FxxA motif of BRC4 occupies the same hydrophobic pocket as I26 from the bacterial RecA linker region that mediates self-oligomerization and filament formation, providing an explanation to the filament-disruptive effect of the BRCs ^36,37^. In vertebrate RAD51 and DMC1, the I26 residue from the linker region of bacterial RecA corresponds to a phenylalanine in a conserved FxxA motif (Figure 1B). The competition between the FxxA motif of the BRC repeats and that of the recombinase linker was further supported by the analysis of the X-ray structure of an archaeal RAD51 heptamer ^38^, the cryo-EM structure of RAD51 nucleofilaments ^39^ and the X-ray structure of *Leishmania* RAD51-ATPase domain in complex with BRC ^40^. The BRC repeat exhibits conserved residues outside of FxxA ^41,42^, including a second hydrophobic patch that sometimes contains phenylalanines and contributes to RAD51 binding, but it is both less conserved and less studied. A seminal study showed that, whereas BRC interacts with monomeric RAD51, TR2 only binds to RAD51 in oligomeric state suggesting an overlap or interaction between interfaces ^31^.

No structure of BRC-DMC1 exists. DMC1 behaves somewhat differently from the other recombinases ^17^ because it has higher propensity to form rings rather than filaments ^43–46^. It was crystallized several times as an octamer, either in its full-length or its N-terminally truncated form. However, even if crystallized as a full-length protein, its 3D structure was defined only from residue I81, suggesting that the N-terminal region from aa 1 to aa 80 is flexible. Cryo-EM structures of DMC1 bound to ssDNA or dsDNA were also reported, in which the N-terminal region is observed and contributes to the oligomerization interface. In all these 3D structures, the invariant phenylalanine F85 from the DMC1 linker region is buried in a hydrophobic pocket of the ATPase domain of the adjacent monomer, thus mediating both self-oligomerization and filament formation.

Other BRC-like FxxA motifs have been spotted in proteins such as RECQL5 ^47^, MCM9 ^48^, FIGNL1 ^49^ that bind RAD51, while a motif similar to PhePP but with a tryptophane in place of phenylalanine was described in the RAD51-binding protein RAD51AP1 and shown to bind DMC1 ^50,51^. However, here again, no structural description of the interfaces between the FxxA and FxPP motifs of these proteins and recombinases is available.

Here we describe the crystal structure of a complex between full-length human DMC1 and the PhePP peptide of BRCA2. This structure revealed that rather than binding to the hydrophobic pocked used by the invariant phenylalanines of the BRC motifs and the self-oligomerization region of the recombinases, the phenylalanine of the PhePP motif uses a distinct, non-overlapping binding interface. Through mutagenesis we determined the importance of the interactions beyond the FxPP consensus. We suggest that our structural analysis also applies to the interaction between RAD51 and BRCA2-CTRD (TR2), and that there are two distinct classes of recombinase-binding motifs, defined by the nature of the residue in the fourth position of the four-residue core motif. The new binding mechanism explains why the P-motifs (FxPP) such as PhePP and CTRD can stabilize recombinase nucleoprotein filament, while the A-motifs (FxxA) of the BRC repeats disrupt and remodel it.

## Results

### BRCA2 PhePP binds DMC1 via a novel interface

In our study describing Brca2Δ12-14 mouse model ^22^, we demonstrated that DMC1 but not RAD51 binds with a detectable affinity to the BRCA2 region that is encoded by exon 14. We sequentially narrowed down the binding domain to the GST-tagged BRCA2 fragments F2 (2351-2479), F2s3 (2379-2433) and F2s6 (2398-2417) that all contain the PhePP motif. A direct interaction between DMC1 and a synthetic peptide corresponding to F2s6 was detected by Isothermal Titration Calorimetry (ITC; Kd = 3.4 µM). We also showed that F2406, the phenylalanine from the PhePP motif, was required for binding. To describe the mechanism of the interaction, we first used AlphaFold to predict the interfaces between PhePP and DMC1, and, for comparison, between BRC4 and DMC1. We obtained high score models for both complexes (interface scores ipTM^52^ above 0.75) (Figure S1A). In the models of DMC1 bound to PhePP, surprisingly, the predictions did not place phenylalanine F2406 into the deep highly conserved hydrophobic pocket that is occupied by F1524 of BRC4 ^41^ and F86 of the linker region of the adjacent monomer ^36,38,44,46^ in the crystal structures of RAD51 (Figure 2A). They systematically positioned F2406 in an alternative hydrophobic pocket located in the ATPase domain, at 15 Å from the one previously described. In the models of the complex between DMC1 and BRC4, the predictions placed phenylalanine F1524 in the hydrophobic pocket identified in RAD51 ^36,38,40^, also occupied by F85 of the linker region of the adjacent monomer in the crystal structures of DMC1 ^47,49^ (Figure 2B).

**Figure 2.**
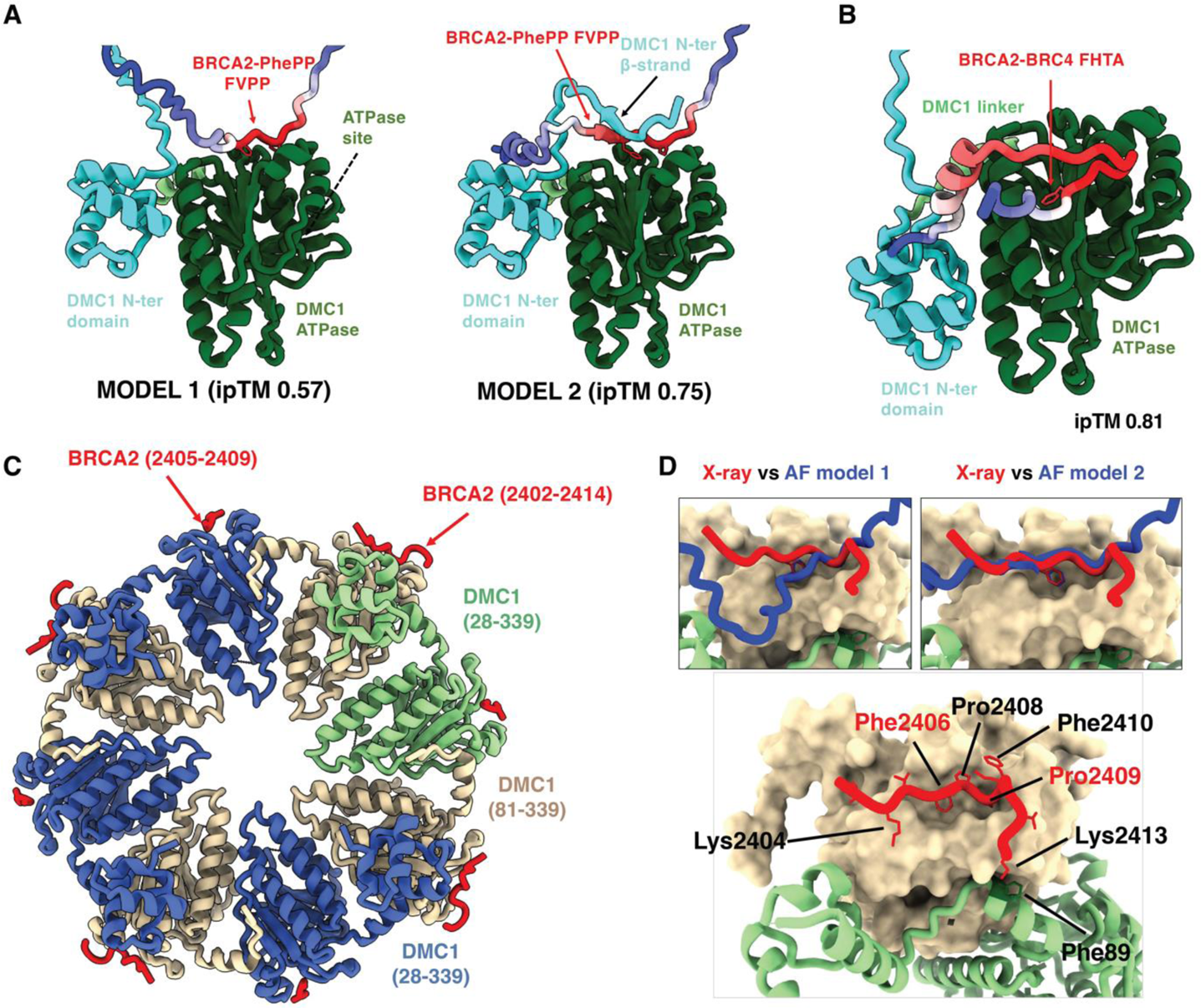
Structure of the DMC1-PhePP complex. (A) AlphaFold models of the PhePP-DMC1 interaction. The BRCA2 peptide is colored as a function of the lDDT score per residue. Side chains of the phenylalanine and proline defining the PhePP motif are displayed in sticks. DMC1 is colored in cyan in its N-terminal region (from M1 to E82), pale green in its linker region (P83 to M97) and dark green in its ATPase domain (V98 to E340). (B) AlphaFold model of the BRC4-DMC1 interaction, displayed as in (A). (C) Crystal structure of the octameric DMC1 (blue and wheat) bound to the BRCA2 peptide F2s6 (red). The blue monomers are observed from Q28 to K339 (one monomer being in pale green), whereas wheat monomers are observed from I81 to K339 (except for a single loop from A272 to either D283 or K285). (D) Focus on the interactions between a wheat DMC1 monomer (surface view) and the adjacent pale green DMC1 monomer (cartoon view) as well as the BRCA2 peptide from P2402 to S2414 (cartoon view). The side chains of the BRCA2 peptide are shown in red sticks. BRCA2 residues that are more than 30% buried upon DMC1 binding are marked (red labels correspond to residues more than 80% buried). The side chain of F89 in the pale green DMC1 monomer, which is located in front of BRCA2 K2413, is displayed in dark green sticks and labeled. In the insets, the positions of the BRCA2 peptides in the X-ray structure (red) and in the AlphaFold models 1 and 2 (blue) are compared by superimposing the interacting DMC1 monomers.

A detailed analysis of the AlphaFold models of DMC1 interacting with PhePP revealed that two different models have medium (ipTM >0.5) or high (ipTM > 0.75) interface scores (Figure 2A; S1A). In both models, the central FVPP motif is buried in the same region of the ATPase domain of DMC1, but in model 2, the N-terminal region of DMC1 (V6-A8) forms an additional β-sheet with the BRCA2 residues preceding the PhePP motif (K2404-F2406). To experimentally test these models, we co-crystallized the recombinant full-length untagged human DMC1 protein (Figure S2) with the synthetic BRCA2 PhePP peptide F2s6 (2398-2417) and determined the structure of this complex by X-ray crystallography at a resolution of 3.46 Å. A dimer consisting of two DMC1-F2s6 complexes was observed in the asymmetric unit, which was assembled into a ring-shaped octamer using the crystallographic symmetries (Figure 2C), as also reported for human DMC1 alone ^44,45^. The dimer comprises monomers defined from F21 (blue and green in Figure 2C) or I81 (wheat in Figure 2C) to K339: one monomer exhibits a homogeneously positioned and folded N-terminal domain (between Q28 and L80), and both monomers show a structured linker region (between I81 to V98) and an ATPase domain (between F99 and K339). Density corresponding to the N-terminal domain is characterized by high B-factors (225-295 A^2^), compared to that of the linker and ATPase regions (150-225 A^2^), revealing that the position of the observed N-terminal domain is still fluctuating in the crystal (Figure S1B). In the ring-shaped octamer, the DMC1 monomers interact mainly through their linker and ATPase regions, as previously reported. The N-terminal domain interacts through a positively charged surface with a negatively charged surface of the ATPase domain of the adjacent monomer (Figure S1C). Two salt-bridges were identified in our crystal structure, between K49 and D204, as well as K74 and E212. Thus, the N-terminal domain contributes to the intermolecular interactions in the DMC1 octamer.

A BRCA2 F2s6 peptide is observed bound to each DMC1 monomer. Its phenylalanine F2406 is buried in a hydrophobic groove on the outer surface of the DMC1 octamer (Figure 2C). The peptide bound to the DMC1 blue monomer is defined from BRCA2 V2405 to P2409, including the FVPP motif, and its density is partial, because this peptide is in a region of contact between two octamers in the crystal. Its position overlaps with that of an equivalent peptide in another octamer (Figure S1D). The peptide bound to the DMC1 wheat colored monomer is defined from BRCA2 P2402 to S2414 (Figure 2D). Its position relative to the wheat monomer is close to that of the peptide bound to the blue monomer (Figure S1E). It is located as predicted by AlphaFold in model 2, which is the best scoring model, but the additional β-sheet formed by the DMC1 N-terminus in this model is not observed in the crystal structure (Figure 2D, insets). At the interface, BRCA2 residues F2406 and P2409 are almost totally buried (more than 80%), whereas residues K2404, P2408, F2410 and K2413 are substantially buried (more than 30%), upon interaction with the wheat colored DMC1 monomer. K2413 is close to F89 of the blue colored DMC1 monomer: their interaction could also help to stabilize the DMC1 oligomer (Figure 2D).

### Binding extends beyond the core FxPP motif and engages the DMC1 N-terminal and linker domain

To support our structural description of the interface between the BRCA2 F2s6 peptide and DMC1 (Figures 3A, B), we proceeded to validate the contribution of the peptide residues buried upon interaction with DMC1. First, we identified positions that are conserved in a multiple alignment of vertebrate BRCA2 sequences around the FVPP motif. This included pairs of positively charged residues on either side of FVPP: R2401··K2404 and K2411·K2413, and the phenylalanine F2410, which is as strictly conserved as F2406 (Figure 3C). Alanine substitutions at the positively charged positions reduced the efficiency of co-precipitation of the GST-tagged PhePP fragments with DMC1 (Figure 3D, S3). Double substitution of the C-terminal pair (K2411A·K2413A) had the strongest effect, abolishing binding as effectively as the F2406A and P2409A substitutions. In contrast, substituting the invariant F2410 for alanine had no effect on the binding. Going back to our crystal structure, we identified that K2404 is close to DMC1 E213, whereas K2413 is close to DMC1 D186 (Figure 3B). Whereas the resolution of our structure is not sufficient to clearly position all the side chains of these residues, our pull-down results supported the presence of salt-bridges between these residues. F2410 is only partially (40% to be compared to 80% in the case of F2406 and 100% in the case of P2409) buried at the interface and seems to play no role in the binding.

**Figure 3.**
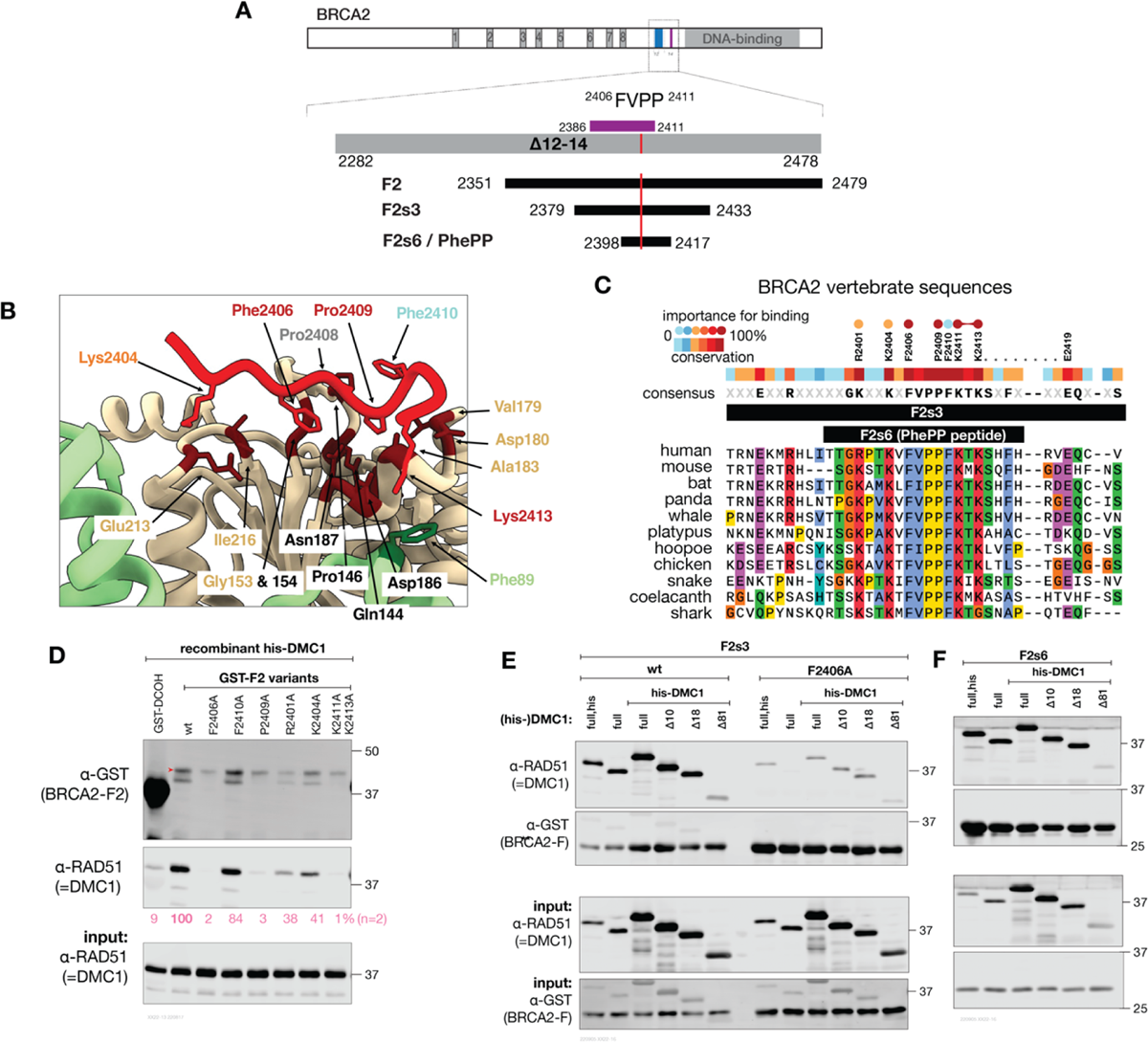
Validation of the DMC1-PhePP binding interface. (A) Schematic of the BRCA2 protein motifs and domain, and the fragments used in this study. The violet bar indicates the PhePP domain as originally defined^14^ (B) Crystal structure of the interface between the BRCA2 PhePP peptide and DMC1, colored as in Figure 2D. In BRCA2, only side chains buried upon binding are displayed, and their label is colored as a function of their contribution to binding, as defined in panels (C,D) (P2408 was not tested). In DMC1, residues that are buried upon binding are brown, their side chains are displayed, and they are labeled in black (conserved in RAD51) or wheat and pale green (not conserved in RAD51). (C) Alignment of human BRCA2 F2s3 (2379-2433) sequence with homologous vertebrate BRCA2 sequences, conservation score and consensus sequence. Conserved positions tested by alanine substitution are displayed above the conservation plot. Circles above indicate the importance of the position for the interaction, as determined in panel (D). (D) GST pull-down between BRCA2-F2 (2351-2479) substitution variants and purified recombinant his-DMC1. The experiment was done twice with the same result, a replica is shown in Figure S3A. Intensity of the bound DMC1 normalized to the BRCA2-F2 lane is shown in pink below the corresponding immunoblot panel (average of the two experiments). (E,F) Interaction between BRCA2-PhePP (fragments F2s3 and F2s6, GST-tagged) and the purified recombinant his-DMC1 truncation variants determined by GST pull-downs. The experiment was done twice with the same result, a replica is shown in Figure S3B.

We also explored the contribution of the interactions with the N-terminal region of DMC1. In AlphaFold model 2, in addition to the DMC1 ATPase domain, the interaction with F2s6 also involved the first 18 amino acids of DMC1, including a small DMC1 motif V6-A8 forming a β-sheet with K2404-F2406 of BRCA2 (Figure 2A). This region of DMC1 is rich in acidic charged side chains (8 out of 18) and is located before the folded DNA binding domain of DMC1 (residues Q28-I81). It is not visible in the crystal structure. To determine if it contributes to binding, we engineered three DMC1 truncation variants, lacking 10, 18 or 81 N-terminal amino acids. Only the largest deletion missing both the acidic region and the N-terminal folded domain had reduced binding to GST-BRCA2 fragment (Figures 3E, F). We concluded that, in solution, the additional β-sheet predicted by AlphaFold is not present at the interface between DMC1 and PhePP. However, the folded domain significantly contributes to binding, even if it is not directly in contact with the BRCA2 peptide. Our crystal structure suggests that a cation-π interaction exists between BRCA2 K2413 and DMC1 linker residue F89, which might be stabilized in the presence of the N-terminal domain (Figure 3B).

### A- and P-motifs and sites

After experimentally establishing the mechanism of interaction between the BRCA2-PhePP motif and DMC1 recombinase, we explored whether these findings extend to other similar motifs and interactions (Figures 4, S4, S5). Since proline in the fourth position (P2409 in PhePP) is essential for binding of FxPP motifs and never occurs in the BRC FxxA motifs, we designate them P- and A-motifs, respectively, and the sites they bind to on the recombinases as P- and A-sites (Figures 1, 4A, B). We used AlphaFold to predict if other P-motifs could bind the P-site (Figures 4C, S4A). For the C-terminal BRCA2 TR2, with a well-established role in RAD51 filament binding and stabilization, AlphaFold predicted binding to the P-interfaces of both RAD51 and DMC1 with medium ipTM scores. For the P-motif of RAD51AP1, the predictions again showed binding to the P-interfaces of the recombinases, despite this motif having tryptophan instead of phenylalanine in the first position. Finally, in FIGNL1, we noticed a previously undescribed P-motif close to its characterized BRC-like motif that binds RAD51 ^49^, and AlphaFold predicted that this motif binds to the P-interfaces of both recombinases.

**Figure 4.**
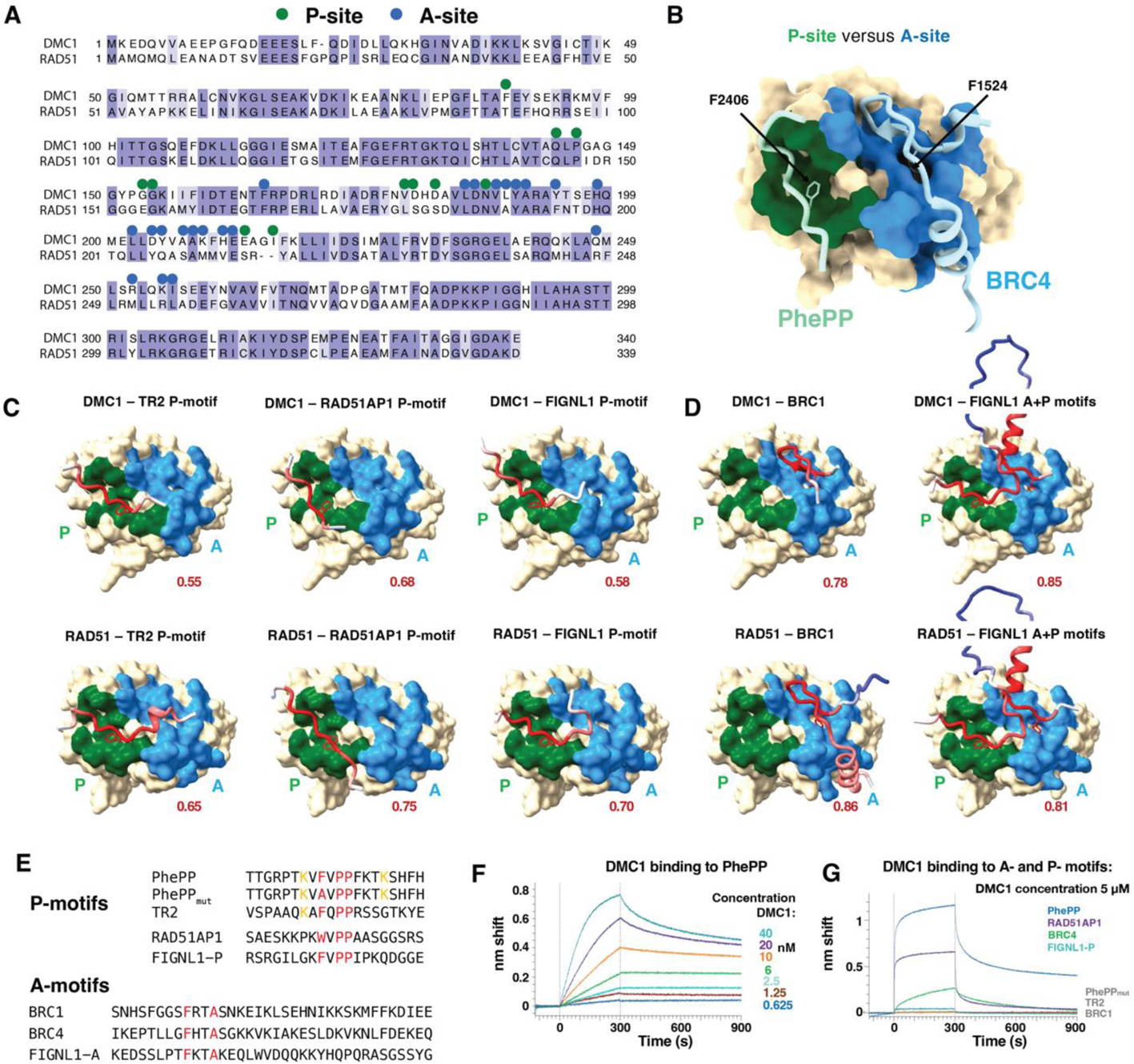
A- and P-recombinase-binding motifs and sites. (A) Alignment of the sequences of human RAD51 and DMC1 recombinases. Green and blue dots correspond to residues buried at more than 30% upon binding of DMC1 to PhePP (P-site) and RAD51 to BRC4 (A-site), respectively. (B) Representation of the surfaces binding to PhePP (P-site; green) and BRC4 (A-site; blue), as defined in (A), onto the 3D structure of RAD51 bound to BRC4 (PDB 1N0W). To display the PhePP peptide bound to the P-site, the crystal structures of DMC1-PhePP and RAD51-BRC4 were superimposed, and the structure of DMC1 was hidden. (C,D) AlphaFold models of the DMC1 and RAD51 complexes with P- and A-motifs from different proteins (see also Fig. S4). The ATPase domain surface of the recombinase is displayed, with the P- and A-sites colored in green and blue, respectively. Peptides containing P- or A-motifs are shown as cartoons, colored based on the AlphaFold IDDT score (red - accurate to white - poorly defined). The phenylalanine side chain of the motifs is shown as sticks. (E) Sequences of P- and A-motif peptides. (F) BioLayer Interferometry experiment performed using immobilized biotinylated PhePP peptide and increasing concentrations of DMC1. The measured apparent Kd is about 1 nM. (G) BioLayer Interferometry experiments revealing that P- and A-motifs bind with different affinities to DMC1 (see also Fig. S5). Using a DMC1 concentration of 5 μM, interaction was detected only in the case of PhePP, RAD51AP1, BRC4 and FIGNL1-P. Apparent affinities for RAD51AP1, BRC4 and FIGNL1 were about 1.6 μM, 4.3 μM, and 16.2 μM, respectively. No binding was detected for PhePP_mut_, TR2 and BRC1.

In parallel, we verified that the A-motif of BRC1, one of BRC repeats with the strongest affinity for both RAD51 ^42^ and DMC1 ^16^, is predicted to bind to the A-site of both RAD51 and DMC1 (Figure 4D, S4B). In FIGNL1, as the A- and P-motifs of FIGNL1 are only separated by 40 residues, we used AlphaFold to calculate models of the whole conserved region containing these motifs and the recombinases. The predictions nicely supported the simultaneous binding of the two motifs to both recombinases.

We then experimentally tested if the identified P-motifs of BRCA2 (PhePP and TR2), RAD51AP1 and FIGNL1, as well as A-motifs of BRCA2 BRC1 and BRC4, bound to DMC1 (Figures 4E-G, S5). Using BioLayer Interferometry (BLI), we found that PhePP is the strongest binder, showing, when immobilized on the biosensor, an affinity of about 1 nM for the recombinase (Figures 4F, S5A). We also detected micromolar interactions between immobilized RAD51AP1 P-motif and DMC1, as well as BRC4 A-motif and DMC1 (Figures 4G, S5). Finally, a weak interaction was observed between FIGNL1 P-motif and DMC1. However, no interaction could be detected when we tested the P-motifs of PhePP mutant F2406A and TR2, as well as the A-motif of BRC1. Altogether, AlphaFold provided interfaces, and BLI experiments gave access to affinities, showing that within the identified P- and A-motifs, predicted as binding to P- and A-sites, respectively, the DMC1 binding energies were distributed on a large range, corresponding to apparent affinities going from 1 nM to 16 µM. Only PhePP variant F2406A, TR2 and BRC1 could not be validated as DMC1 binders in our conditions.

### PhePP binding stabilizes DMC1-DNA nucleoprotein filaments

Our crystal structure revealed that PhePP binds to oligomeric DMC1 ATPase domain, and, together with our pull-down experiments, strongly suggested that PhePP contacts two adjacent DMC1 monomers (Figure 5A). Mutating DMC1 phenylalanine F85 is predicted to impair DMC1 oligomerization, as experimentally demonstrated in the case of the RAD51-TR2 complex ^31^, which should then decrease PhePP binding. On the other hand, BRC4 is predicted to only bind to DMC1 monomers (Figure 5A). To experimentally support these predictions, we tested binding of the GST-fused peptides PhePP, PhePP mutant F2406A and TR2 to DMC1 either wild-type or F85A. We observed that the F85A substitution substantially reduced DMC1 binding to PhePP (Figures 5B, S6), but had an opposite effect on binding to BRC4, as expected. The impact of F85A on TR2 binding was similar to its effect on PhePP binding, suggesting that both P-motifs preferentially bind oligomeric recombinase. This is in perfect agreement with the conclusions of a previous study that concluded that TR2 can only bind RAD51 oligomers and not monomers ^31^.

**Figure 5.**
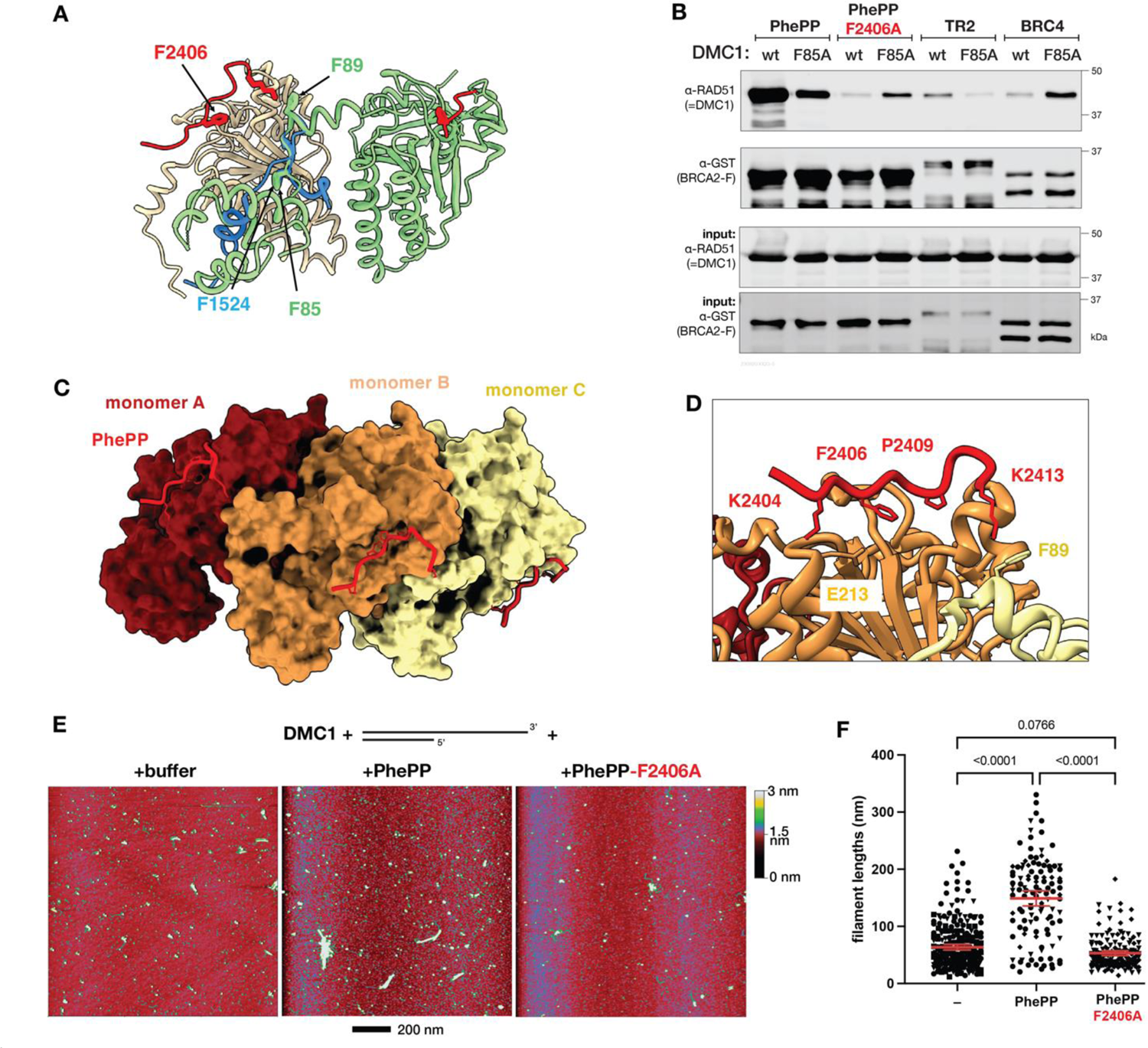
PhePP binding stabilizes DMC1 oligomers. (A) Superposition of the crystal structure of a dimer of DMC1-PhePP onto the crystal structure of RAD51 ATPase domain bound to BRC4. The two DMC1 monomers are in wheat and green, as in Figure 1C. The PhePP peptides are in red. RAD51 is not displayed and the BRC4 peptide is in blue. In DMC1 green monomer, the side chains of F85 at the interface with the wheat monomer and F89 at the interface with PhePP are shown in sticks. In PhePP, the side chains of two residues important for binding to DMC1, i.e. F2406 and K2413 (interacting with F89), are also represented. In BRC4, only the side chain of F1524, essential for binding RAD51, is displayed. (B) GST pull-down assay with GST-tagged BRCA2 peptides and either wild type DMC1 or its F85A variant (replicated in Figure S6). (C) Docking of three PhePP peptides onto the cryo-EM structure of the DMC1-ssDNA filament (PDB 7C9C). Each peptide was positioned by superimposing the DMC1 filament monomer (either A, B or C) onto the crystal structure of DMC1 (81-339) bound to PhePP, as shown in Figure S7. DMC1 (81-339) structures were subsequently hidden. The ssDNA binds on the opposite side of the filament, as compared to PhePP, it is thus not visible in this view. (D) Zoom from (C). The side chains of PhePP important for binding to DMC1 are displayed in sticks. The side chains of the DMC1 residues in front of the two lysines are displayed. The same side chains were identified in front of the two lysines in the crystal structure of DMC1-PhePP. (E) Representative images of the SFM scans of the complexes formed by DMC1 on a DNA molecule corresponding to resected DSB end, in the presence of the F2s3 (PhePP) peptide, its F2406A variant or storage buffer as negative control. (F) Quantification of the filament lengths in the SFM data shown in panel (C). Symbol shapes correspond to independent experiments (n=), three (–, PhePP) or two (PhePP F2406A). Red lines indicate mean and 95% confidence intervals. Significance was determined by ANOVA with Kruskal-Wallis test.

We then asked if this interdependence between DMC1 oligomerization and PhePP binding efficiency could stabilize the DMC1 nucleoprotein filament, as has been reported for TR2-DMC1. We first modeled a ternary complex between DMC1, ssDNA and PhePP, by fitting the crystal structure of DMC1 bound to PhePP onto the cryo-EM structure of DMC1 within the presynaptic filament ^53,54^(Figure S7). Analysis of the resulting model showed that there is no steric clash between the peptide and the filament (Figure 5C). The local environment of the PhePP residues is similar in the crystal structure and in the DMC1-PhePP-ssDNA model (Figure 5D). The P-motif is buried in the hydrophobic cavity observed in our crystal structure, K2404 is still in front of E213 in DMC1 ATPase domain, and even K2413 is close to F89, suggesting that it is still able to make a cation-π interaction with the linker domain of the adjacent DMC1 monomer. To experimentally test the impact of PhePP on the formation of the DMC1 nucleoprotein filament, we observed by scanning force microscopy (SFM) DMC1 filaments formed on tailed DNA substrates (505 bp dsDNA with 289 nt 3’-overhang). We found that in the presence of PhePP, DMC1 filaments were about two times longer than in its absence or when PhePP-F2406A variant was added (Figures 5E, F). This significant difference together with our structural analysis and pull-down results revealed that P-motif binding is not only compatible with the recombinase oligomerization into a nucleoprotein filament, but it actually depends on it and stabilizes it.

## Discussion

The essential role of BRCA2 in somatic and meiotic HR stems from its interactions with the RAD51 and DMC1 recombinases mediated by short motifs in the central and C-terminal regions of BRCA2 (Figure 1). The motifs are very similar: the core sequence is four amino acids long with an invariant phenylalanine in the first position and a highly constrained or invariant residue in the fourth (alanine or proline). The recombinases are also very similar in sequence and interactions with BRCA2, and nearly identical structurally ^44^. Yet in meiosis, where both of them are present, they behave differently. To which extent these functional differences are intrinsic or are regulated by accessory proteins like BRCA2 was not clear. We recently described a mouse strain with a deletion in BRCA2 that resulted in crucial separation of function: DMC1 localization to meiotic DSBs in oocytes was abolished, but RAD51 was not. Here we provide a molecular mechanistic explanation for that mouse phenotype. We determined the crystal structure of DMC1 in complex with the PhePP motif from the deleted region, analyzed the interaction computationally and biochemically and established its effect on DMC1 activity. The structure led to two key findings. The first is that despite the similarity in sequence, the phenylalanine-containing motifs bind to different sites. We classified the motifs and the corresponding interfaces into A-(FxxA) and P-(FxxP) and propose that our findings with PhePP-DMC1 extend to other P-motifs, and in particular the recently identified P-motif in TR2. Second, we provide further evidence and molecular explanation to the opposite effects of the A- and P-motifs on the ability of the recombinase to oligomerize into filaments.

The P-site on the DMC1 monomer ATPase domain in our structure contacts BRCA2 residues T2403-K2413 through a surface of about 500 Å^2^, which is half the size of the A-interface in the BRC4-RAD51 structure ^36^. Two BRCA2 residues, F2406 and P2409, contribute most to the P-interface (buried surface area per residue > 100 Å^2^), and their mutation to alanine impairs binding. The two lysines K2404 and K2413 are also largely buried upon binding (> 80 Å^2^). K2413 is close to the phenylalanine F89 from the linker region of the adjacent DMC1 monomer. Substitution of K2413 for alanine strongly decreases binding to DMC1, suggesting that BRCA2 K2413 and DMC1 F89 establish a cation-π interaction, so that the BRCA2 PhePP peptide simultaneously interacts with two adjacent DMC1 monomers. K2404 is close to the N-terminal domain of the adjacent DMC1 monomer. Their interface is poorly defined in our crystal structure. However, we found that K2404A substitution substantially decreases DMC1 binding and PhePP poorly binds to DMC1 lacking first 81 amino acids, which supports the existence of this interface. Also, previous crystallographic studies of octameric full-length DMC1 did not show electron density corresponding to the N-terminal region of DMC1 ^44,45^, and in our study, we did not find any crystal contact restraining the position of the observed N-terminal domain, suggesting that PhePP binding stabilized the position of this domain and its interaction with the adjacent DMC1 ATPase domain. Consistently, we experimentally found that the dependence on oligomerization is reciprocal: PhePP not only binds to the oligomeric form of DMC1, interacts less with DMC1 mutant F85A which has a defective oligomerization interface, but also stabilizes DMC1 oligomers in the functional filament configuration (Figure 5C, D). In our model of the DMC1-ssDNA-PhePP complex, PhePP binds to DMC1 ATPase domain through the same interface core as observed in the DMC1-PhePP crystal structure. Moreover, the interaction between BRCA2 K2413 and DMC1 F89 of the adjacent monomer is conserved in the filament. We conclude that the P-site extends beyond the DMC1 ATPase domain of one monomer to the linker of the adjacent monomer, with the N-terminal domain being able to indirectly contribute to this interface, which altogether favors the formation of the DMC1 nucleofilament and stabilize it.

Our study benefited from the power of AlphaFold modeling. The prediction of a novel interaction mechanism for DMC1 and PhePP was unexpected, and we were unable to trace its origin to any known recombinase structure. It was fully corroborated experimentally by the crystal structure (Figure 2), mutagenesis (Figure 3) and functional assays (Figure 5). Therefore, based on additional AlphaFold modelling, sequence analysis and previously published data including our own, we hypothesized that other P-motifs will interact with recombinases via the same interface and with similar functional consequences. Among these, the P-motif with the TR2 domain is most notable. Our findings on PhePP-DMC1 are fully consistent with and suggest a molecular explanation to the seminal observations on RAD51-TR2 published by Davies and Pellegrini: preferential binding to oligomeric and reduced binding to monomeric form of recombinase and the stabilizing effect, which can counteract the disruptive effect of BRC. It is also supported by multiple studies that showed that the C-terminal domain of BRCA2 promotes RAD51 function and replication fork protection in cells. These biochemical and cell biological studies focused on phosphorylation of S3291 and the adjacent P-motif positioned next to it has only recently attracted attention ^30^. As S3291 substitutions caused only mild phenotypes in cells and animals, it may have a modulatory effect, while the adjacent P-motif defines the function. The role of the conserved region of FIGNL1 L292-K346, containing both A-and P-motifs that might simultaneously bind to recombinases, is also intriguing. The phenylalanine of the A-motif is essential for binding RAD51, but the P-motif is also involved in this interaction ^49^. We consistently observed only a weak binding of the P-motif alone to DMC1. All this data suggests that the A- and P-motifs of FIGNL1 simultaneously bind to the contiguous A- and P-sites in RAD51 and DMC1, which might promote the dissociation of the recombinases from nucleoprotein complexes at specific stages of HR.

The definition of two classes of recombinase-binding motifs, which bind to distinct but contiguous A- and P-sites, calls for re-evaluation of the previous data on recombinases and proteins that bind them and points to new experiments to explore them. The large sequence diversity of the A- and P-motifs, outside of their core FxxA and FxxP residues, suggests a variability in function (Figure 4E). The important lysine residues identified in PhePP are only partially conserved in TR2 and not conserved in the P-motifs of RAD51AP1 and FIGNL1, even if all these motifs are highly positively charged. How the distribution of the positively charged residues regulates binding of P-motifs to two adjacent recombinase monomers (π-cation interaction, involvement of the linker and the N-terminal domain) is still unclear. Thus, our work provides a molecular explanation to DMC1 disruption in Brca2Δ12-14 mouse and to RAD51- and replication fork-stabilizing effects described for the BRCA2 TR2/CTRD/CTRB and opens a new avenue of mechanistic questions on the disruptive versus stabilizing effect of A- and P-motifs outside of BRCA2.

## Materials and Methods

### Protein Expression and Purification

DMC1 was purified exactly as described in our paper describing the *Brca2^Δ^*^12–14^ mouse model ^22^. Construct pAZ379 for bacterial expression of human his-TEV-DMC1 was engineered by subcloning DMC1 coding sequence into pETM-11 vector using Gibson assembly. Rosetta2 DE3 pLysS *E. coli* strain was transformed with the construct and plated on selective (50 µg/mL kanamycin, 30 µg/mL chloramphenicol) LB-agar plates. Ten colonies were used to inoculate 200 mL selective LB media, the culture was grown overnight at 37 °C with shaking and used to inoculate 9 L selective LB media. The 9 L culture was grown at 37 °C till OD_600_ reached 0.6-0.8, expression was induced by adding IPTG to 0.2 mM final concentration. Cells were cultured for another 3 h at 37 °C, collected by centrifugation and frozen. The pellet was thawed in two volumes of lysis buffer (3M NaCl, 100mM Tris pH7,5, 10% Glycerol, 0.5mM EDTA, 5 mM β-mercaptoethanol, protease inhibitors (Roche)) and sonicated on ice (10 pulses of 10 seconds). The lysate was incubated overnight at 4 °C on a rotator, and then cleared by centrifugation at 22000 rcf for 45 minutes at 4 °C. Cleared supernatant was supplemented with 5 mM imidazole and incubated with washed (500 mM NaCl, 25 mM Tris pH 8, 10% Glycerol, 0.5 mM EDTA, 1 mM DTT) Ni-NTA beads for 1 h at 4 °C on a rotator. The beads were washed with 20 mM and 50 mM imidazole (two times 10 mL for each wash). Bound protein was eluted with 400 mM imidazole, fractions analyzed by electrophoresis and pooled, supplemented with TEV protease and dialyzed overnight against 2 L of 150mM KCl, 10% Glycerol, 50mM Tris pH 8, 5mM β-mercaptoethanol. Protein was further purified using 5 mL HiTrap heparine and CaptoQ columns.

### Tailed DNA substrate

DNA with a 3’ ss overhang was made as follows: using the URA3 gene from Saccharomyces cerevisiae as template DNA, an 810-bp polymerase chain reaction (PCR) fragment was produced using primer URA3 (GAAGGAAGAACGAAGGAAGGAGC) which was 5’ phosphorylated and primer BIO 5’ (TTTCCCGGGGGGCCCGGGTTCTATACTGTTGACCC) which was 5’ biotinylated. The PCR product was purified on a GFX^TM^ column (GE Healthcare) and incubated with 1 mg of streptavidin-coated magnetic beads (Dynabeads M-270, Invitrogen) for 2 h at 37^0^ C in 400 μl of BW buffer (5 mM Tris–HCl pH 7.5, 0.5 mM EDTA, 1 M NaCl). Using a magnetic particle concentrator, the immobilized DNA was washed two times with 100 μl of BW buffer.

Subsequently, the DNA was washed once with 100 μl of 1x λ exonuclease buffer (NEB). The DNA bound to the beads was made ss by λ exonuclease digestion of the strand with 5’ phosphate (5 U/mg DNA, NEB). The reaction was carried out at 37^0^ C for 1 h. Immobilized ssDNA was washed with 100 μl of BW buffer. Subsequently, the DNA was washed once with 100 μl of 1x Sequenase buffer (USB) and hybridized with oligonucleotide primer DR3 (GGTCCCAAAATTTGTTTACTAAAA), which anneals at position 290–313 along the ss PCR product. Primer DR3 was extended for 30 min at 37^0^ C in a reaction containing Sequenase^TM^ DNA polymerase Version 2.0 (USB) in 100 μl of 1x Sequenase buffer supplemented with 200 mM dNTPs. The resulting DNA with 505 bp ds region and 289-nt 3’ overhang was washed with 100 μl of BW buffer, released from streptavidin-coated magnetic beads by digestion with Sma I (ROCHE) and purified on a GFX^TM^ column.

### Scanning force microscopy

DMC1 (30 nM) was incubated in buffer containing: 30 mM HEPES-KOH pH 7.2, 150 mM NaCl, 1 mM CaCl_2_ and 1 mM DTT for 10 min and deposited onto freshly cleaved mica. After 10 s the mica surface was washed with milli Q water and dried with a stream of filtered air. DMC1–tailed DNA complexes were formed in 20-μl reactions containing 3.75 μM DNA, 2.5 μM DMC1, 30 mM HEPES-KOH pH 7.2, 150 mM NaCl, 1 mM CaCl_2_ 1mM ATP and 1 mM DTT. Reaction mixtures were incubated at 37 °C for 10 min, diluted 15-fold in deposition buffer (10 mM HEPES-KOH (pH 7.5) and 10 mM MgCl_2_) and deposited on freshly cleaved mica. After 10 s, the mica was washed with water and dried in a stream of filtered air. When the effect of the presence of PhePP on filament formation of DMC1 was studied, DMC1 was first preincubated with PhePP or PhePP-F2406A on ice for 10 min (6-fold molar excess of peptide) and then DMC1–tailed DNA complexes were formed in 20-μl reactions containing 1 μM DMC1, 3.75 μM DNA, 30 mM HEPES-KOH pH 7.2, 150 mM NaCl, 1 mM CaCl_2_ 1mM ATP and 1 mM DTT. Reaction mixtures were incubated at 37 °C for 10 min, diluted 10-fold in deposition buffer (10 mM HEPES-KOH (pH 7.5) and 10 mM MgCl_2_) and deposited on freshly cleaved mica. After 10 s, the mica was washed with water and dried in a stream of filtered air. RAD51–tailed DNA complexes were formed in 20-μl reactions containing 3.75 μM DNA, 1.25 μM RAD51, 50 mM Tris-HCl (pH 7.5), 2 mM CaCl_2_, 1 mM ATP, 1 mM DTT and 30 mM KCl. Reaction mixtures were incubated at 37 °C for 10 min, diluted 15-fold in deposition buffer (10 mM HEPES-KOH (pH 7.5) and 10 mM MgCl_2_) and deposited on freshly cleaved mica. After 10 s, the mica was washed with water and dried in a stream of filtered air. Images were obtained on a NanoScope IV SFM (Digital Instruments; Santa Barbara, CA) operating in tapping mode in air with a type J scanner. Silicon Nanotips were from AppNano (Santa Clara, CA).

### Crystallization of DMC1/PhePP complex and structure determination

Crystallization assays of DMC1/PhePP complexes were performed at 293 K in the High Throughput Crystallization (HTX) laboratory (EMBL Grenoble). The complex was prepared by mixing the protein DMC1 with the PhePP peptide at a molar ratio of 1:2.5. The final protein concentration was at 7.4 mg/ml. Crystallization screens were performed by sitting drop vapor diffusion, mixing 100 nl of sample with 100 nl of reservoir solution. Several crystallization conditions were identified within several days and the crystals were grown for a month. They were prepared for X-ray diffraction experiments using the CrystalDirect harvesting and processing robot ^55^. The crystals were flash-frozen in liquid nitrogen without cryoprotection. The best crystals were obtained using a crystallization solution of 0.1 M MES at pH 6 or 2.4 M of sodium formate at pH 6, and diffracted up to 3.46 Å of resolution. X-ray diffraction data were collected on the MASSIF-1 beamline at ESRF synchrotron (Grenoble). Diffraction data were indexed, integrated and scaled using XDS ^56^. The crystal structure was determined by molecular replacement with MOLREP ^57^ using the human DMC1 ATPase domain crystal structure (PDB code 6R3P, Dunce, J.M. & Davies, O.R., unpublished) and the AlphaFold2 predicted structure of the N-terminal domain as search models. Refinement was performed with BUSTER-2.10.4 ^58^ using TLS groups, NCS restraints and target geometrical LSSR ^59^ restraints from the higher resolution structure (PDB code 6R3P). Inspection of the density map and manual rebuilding were performed using COOT ^60^. Molecular graphic images were generated using ChimeraX ^61^. In the final model, DMC1 chain A was modeled from Q28 to V62, from L65 to T271 and K286 to K339, the DMC1 chain B was modeled from I81 to T271 and P284 to K339, the PhePP peptide C was modeled from P2402 to S2414, and the PhePP peptide D was modeled from V2405 to P2409. Crystallographic data and refinement parameters are summarized in Table S1. The crystal structure was deposited at the Protein Data Bank as 8QQE.

### GST pull-down assay

GST pull-down assays were performed exactly as described in our paper describing the *Brca2^Δ^*^12–14^ mouse model^22^. Bacterial expression constructs for expression of his-GST-TEV-tagged BRCA2 fragments and their amino acid substitution variants were engineered using Gibson assembly in pETM-30 vector and sequence-verified. Constructs were transformed into Rosetta2 DE3 pLysS *E. coli* expression strain. Two mL of selective LB (50 µg/mL kanamycin, 30 µg/mL chloramphenicol) media was inoculated, the culture was grown overnight at 37 °C with shaking, diluted to 10 mL, further incubated till OD_600_ reached 0.6-0.8, induced with 0.2 mM IPTG, grown for additional 3 h at 37°. Cells were collected by centrifugation, pellet was resuspended in 1 mL NETT+DP buffer (NaCl 100 mM, Tris-HCl pH 7.5 50 mM, EDTA pH 8 5 mM, Triton X100 0.5%, freshly supplemented with protease inhibitors (Roche), 1 mM DTT, 1 mM PMSF) and sonicated (10× 5 sec on, 10 sec off). Lysate was transferred to Eppendorf minicentrifuge tubes and cleared by centrifugation in (30 min at 4 °C). Supernatant was mixed with 20 µL GSH-Sepharose beads (GE Healthcare 17-5132-01) and incubated overnight at 4 °C. Beads were collected by centrifugation (500 rcf, 2 min, 4 °C), washed with NETT+ buffer ((NaCl 100 mM, Tris-HCl pH 7.5 50 mM, EDTA pH 8 5 mM, Triton X100 0.5%, freshly supplemented with protease inhibitors (Roche)), and incubated with DMC1 protein solution (∼2 µg in 1 mL NETT+) for 1.5 h at 4 °C. Prior to incubation, bead suspension was vortexed, a 40 µL aliquot was collected as input, mixed with 12 µL 5× sample buffer (50% glycerol, 250 mM Tris HCl pH 6.8, 10% SDS, 0.5% bromophenol blue, 0.5 M β-mercaptoethanol) and denature for 5 min at 95 °C. After incubation with DMC1, beads were washed three times with NETT+ buffer and incubated with 25 µL of Laemmli sample buffer for 5 min at 95 °C to elute bound proteins. Samples (5 µL of eluate) were run on a 13% tris-glycine acrylamide gel, transferred to PVDF membrane, blocked (5% skim milk in PBS-T (PBS with 0.05% Tween-20)) and immunodetected with a mixture of rabbit anti-RAD51 pAb (1:20000 ^62^) and mouse anti-GST mAb (1:5000, B-14 Santa Cruz, sc-138) antibodies overnight. After washes, the membranes were incubated with fluorescently-labelled secondary antibodies (anti-mouse CF680 (Sigma #SAB460199), anti-rabbit CF770 (Sigma #SAB460215)), washed (5 × 5 min PBS-T) and scanned using Odyssey CLx imaging system (LI-COR).

### ATPase activity assay

Non-radioactive colorimetric malachite green ATPase activity assay was performed in 96-well mictotiter plates in triplicates. Prior to experiment, all glassware was thoroughly rinsed with ultrapure milli-Q water to remove all traces of phosphate. Solution AM was prepared by adding to 25 mL milli-Q water 17.25 mL fuming HCl, then slowly dissolving 2.1 g ammonium heptamolybdate * 4H2O (Sigma-Aldrich A7302) and adjusting the volume to 50 mL with milli-Q water. Solution MG was prepared by dissolving 67.5 mg malachite green (Sigma-Aldrich M6880) in 100 mL and then adjusting to 150 mL milli-Q water. Solution MGAM was prepared by mixing 150 mL MG and 50 mL AM for at least 30 min and filtering through 0.22 µm PES or cellulose acetate filter. The solution was stored at 4 °C and used in multiple experiments. Before the experiment an aliquot of MGAM solution was complemented with Tween-20 (0.01% final). The reaction containing 1 µM DMC1 or RAD51 in 20 µL reaction buffer (50 mM Tris pH 7.5, 150 mM NaCl, 1.5% glycerol, 2 mM DTT, 2 mM MgCl_2_, 100 ng/µL acetylated BSA, 50 µM (nucleotides) phiX174 ssDNA) were pipetted into wells. 5 µL of 1 M ATP solution was added to different wells every 15 min for the duration of the time course. The reaction was stopped by adding 100 µL MGAM-Tween solution. Two minutes later 12 µL of 34% sodium citrate (Sigma-Aldrich 71405) solution were added. Absorbance at 620 nm was read 20-40 minutes later. Standard phosphate solutions were in concentration range 0-160 µM.

### AlphaFold calculations

Initial structure predictions were obtained using AlphaFold2 as implemented in the ColabFold version 1.3.0. Multiple sequence alignments were obtained from the MMSeqs2 MSA server in the mode: “UniRef+ Environmental”. For ColabFold, all default parameters were kept, notably model_type: “AlphaFold2-ptm” and rank_by: “plddt”, except for num_recycles that was set to 18. No *Amber* relaxation or any manual interventions were applied to the models. All predictions were repeated using a similar protocol on the Integrative Bioinformatics platform of I2BC (https://bioi2.i2bc.paris-saclay.fr). For each complex, a series of five models were calculated and the models were analyzed using the delivered heatmap and lDDT plots, as well as the pTM- and ipTM scores ^52,63^.

### BioLayer Interferometry (BLI)

Biomolecular interactions between DMC1 and biotinylated peptides BRCA2 PhePP: Biotin-GSG-TTGRPTKVFVPPFKTKSHFH, BRCA2 PhePPmut: Biotin-GSG-TTGRPTKVAVPPFKTKSHFH, BRCA2 TR2: Biotin-GSG-VSPAAQKAFQPPRSSGTKYE, FiGNL1 P-motif: Biotin-GSG-RSRGILGKFVPPIPKQDGGE; RAD51AP1 P-motif: Biotin-GSG-SAESKKPKWVPPAASGGSRS, BRCA2 BRC1: Biotin-GSG-SNHSFGGSFRTASNKEIKLSEHNIKKSKMFFKDIEE; BRCA2 BRC4: Biotin-GSG-IKEPTLLGFHTASGKKVKIAKESLDKVKNLFDEKEQ (synthesized by Proteogenix and Genecust) have been measured by BLI using an Octet RED96 instrument (FortéBio) and Streptavidin (SA) biosensors. The measurements were made at 25 °C. The biosensors were hydrated for 20 min in 25 mM Tris, pH 7.5, 100 mM NaCl, 5mM bMe buffer (the same buffer as for the DMC1 protein). For the binding assays, the biotinylated peptides were immobilized on Streptavidin sensors by immersing the sensors with 0.25 µg/mL peptide solutions. No binding of DMC1 was observed in the binding assays without loaded biotin peptides. The SA biosensors were washed with the buffer before the incubation (300 sec) with DMC1 at different concentrations to record the association processes. The kinetic assays were performed at different ranges of concentrations of DMC1, using two-fold serial dilution, as indicated. Dissociation processes were measured by immersing the sensors in the buffer for 600 sec. 5 cycles of 10 seconds incubation in the regeneration buffer (1 M NaCl) followed by 10 seconds incubation in the assay buffer (25 mM Tris, 100 mM NaCl, 5mM bMe, 0.05% Tween) were made before each baseline for every kinetic experiment. The KD were measured in the steady-state mode.

## Acknowledgements

S.M. and S.Z.J. were supported by funding from the French Infrastructure for Integrated Structural Biology [https://www.structuralbiology.eu/networks/frisbi; ANR-10-INSB-05-01] and the European Community H2020 Programme under the project iNEXT Discovery (Grant No 871037). S.M. also acknowledges funding by the Institut National de la Santé et de la Recherche Médicale (INSERM). We thank Chloé Quignot for implementing the AlphaFold-multimer at Institute of Integrated Biology of the Cell (I2BC). We also thank Florine Dupeux and all the HTX Lab staff for the crystallogenesis and crystallography experiments, and Magali Nicaise-Aumont on the Macromolecular Interaction Platform of I2BC for access and help when working on the BLI instrument. We acknowledge ESRF and SOLEIL for the provision of synchrotron radiation facilities. This study was supported by the Oncode Institute, which is partly financed by the Dutch Cancer Society (KWF). We thank the Josephine Nefkens Cancer Program for infrastructure support.

## Supplementary Figures

**Supplementary Figure S1 (related to Figure 2).**
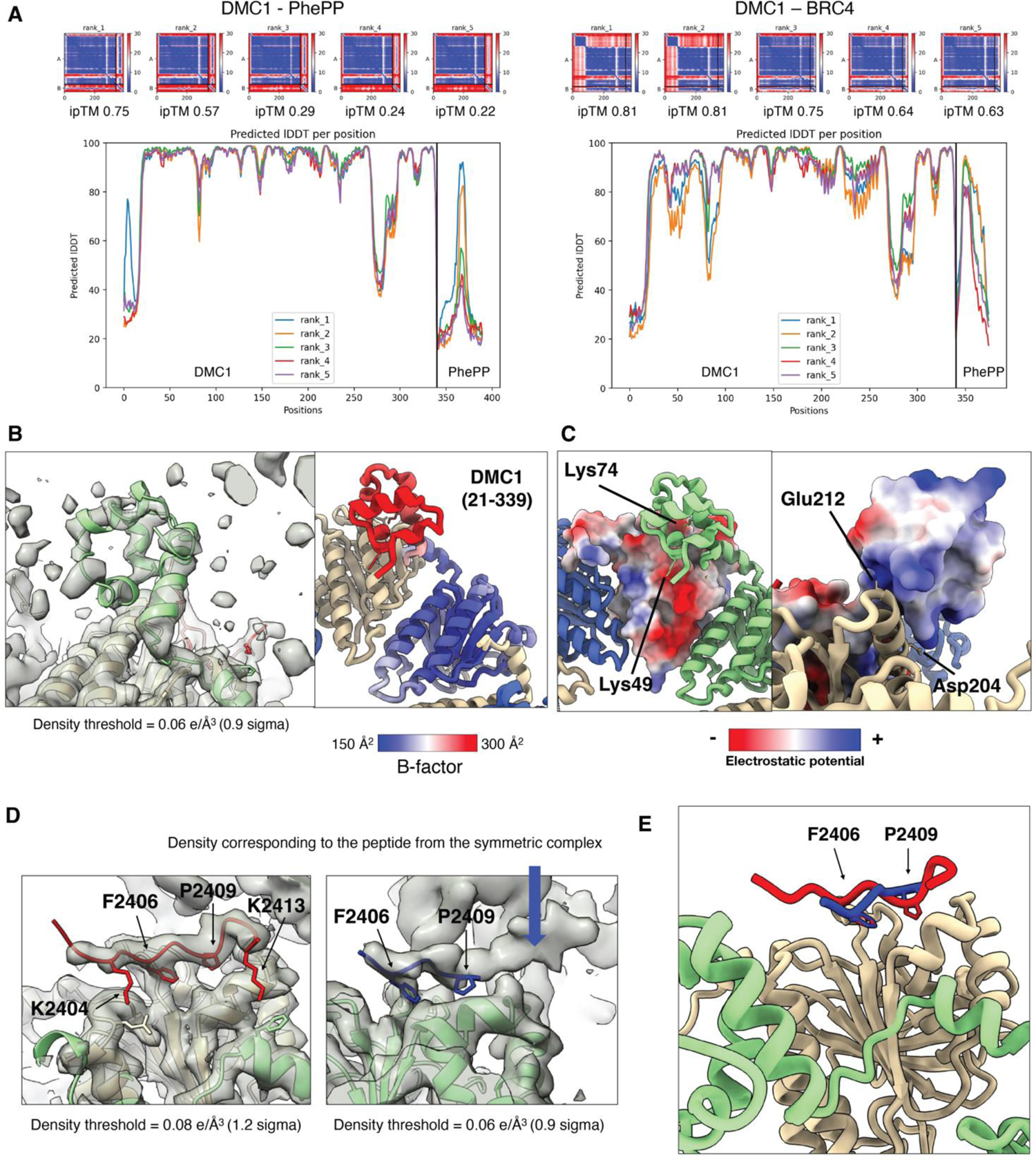
Structural analysis of the DMC1-PhePP complex. (A) AlphaFold plots scoring the 5 models calculated for DMC1-PhePP and DMC1-BRC4, respectively. In the upper panels, the heat maps show, for each of the 5 calculated models, the predicted relative position error (in Å) calculated between all pairs of residues (DMC1 and PhePP residues are numbered as 1-340 and 341-390, respectively; the color scale goes from blue - no error - to red - error above 30 Å). In the lower panels, the lDDT score is plotted as a function of the residue number. (B) DMC1 N-terminal domain observed in the electron density of the complex. In the left panel, the sigmaA weighted 2Fo-Fc electron density map is shown in grey, and the structure of the DMC1-PhePP complex is displayed as in Figure 2C. In the right panel, a cartoon representation of a monomer of DMC1 observed between residues Q28 and K339 is colored as a function of the B-factors, all the other monomers being colored as in Fig. 2C. (C) Electrostatic potential at the interface between the N-terminal domain and the ATPase domain of the adjacent monomer. Left: surface representation of a monomer of DMC1 observed between residues 81 and 339, colored as a function of the electrostatic potential, all the other monomers being colored as in Figure 2C. The side chains of the two lysines from the green monomer located in front of negatively charged residues are displayed as sticks and labeled. Right: surface representation of a monomer of DMC1 observed between residues 28 and 339, colored as a function of the electrostatic potential, all the other monomers being colored as in Figure 2C. The side chains of the two negatively charged residues from the wheat monomer located in front of the lysines indicated on the left panel are displayed as sticks and labeled. (D) PhePP peptides (F2s6 construct) observed in the electron density of the complex. The sigmaA weighted 2Fo-Fc electron density map is shown in grey, and the structure of the DMC1-PhePP complex is colored as in panel (E). Side chains of the most buried residues of the PhePP peptides are represented as sticks and labeled. In the left panel, the side chains of DMC1 E213 (wheat) and F89 (green) are also displayed in sticks. (E) Superposition of the structures of the BRCA2 peptides bound to the wheat and green monomers, respectively. The peptide in red is bound to the wheat monomer. To position the peptide in blue, the green monomer was fitted onto the wheat monomer and its structure was hidden.

**Supplementary Figure S2.**
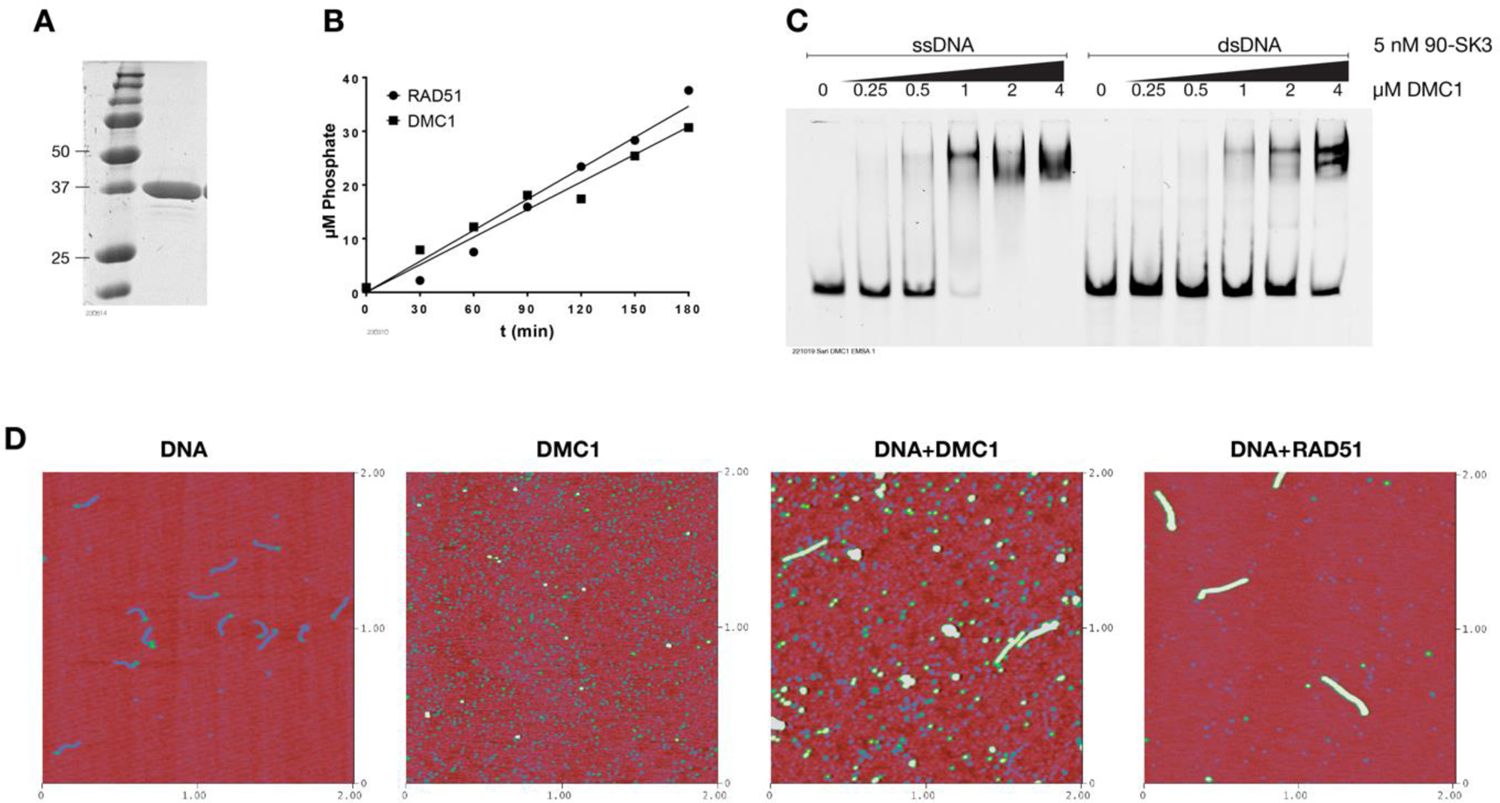
Functional activity of purified DMC1. (A) Coomassie brilliant blue staining of purified DMC1 analyzed by SDS-PAGE. (B) ATPase activity of purified DMC1 and RAD51 proteins. (C) Binding of DMC1 to single- and double-stranded DNA oligos, fluorescently labelled analyzed by EMSA. (D) SFM analysis of tailed DNA substrate, DMC1, and filaments formed by DMC1 and RAD51 on tailed DNA substrate.

**Supplementary Figure S3.**
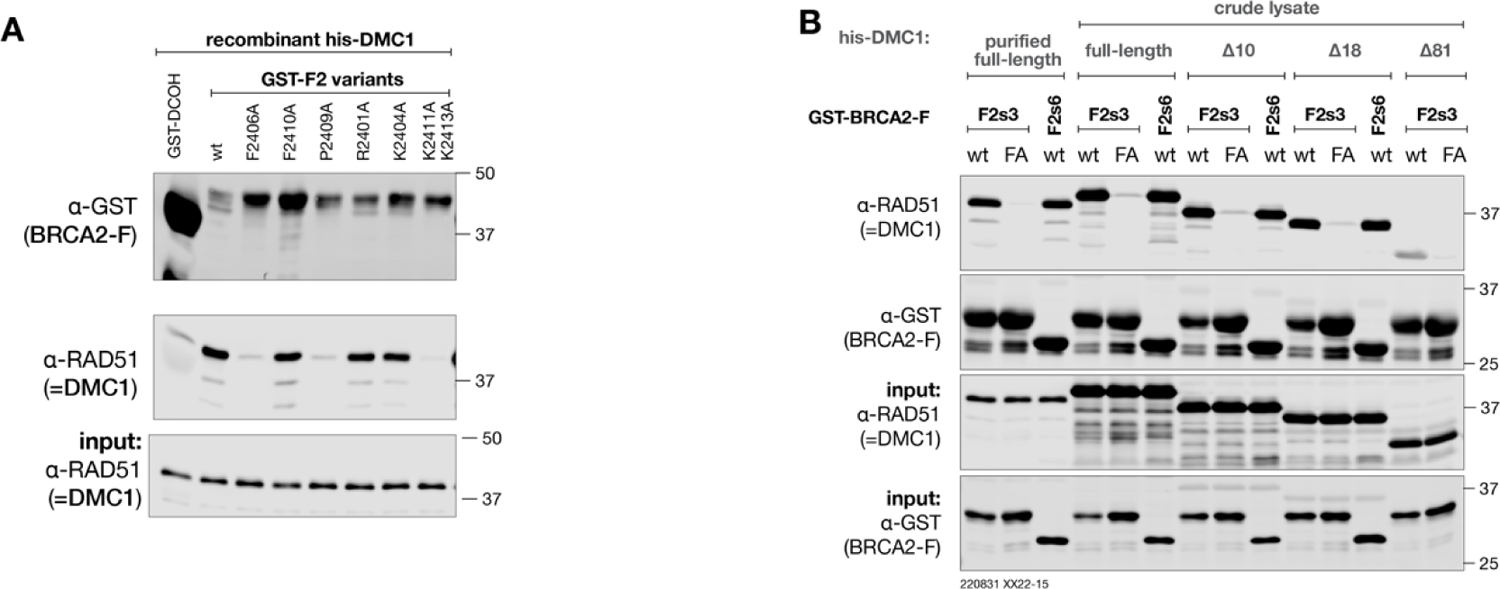
Replicates of the experiment shown in Figure 3 D,E.

**Supplementary Figure S4 (related to Figure 4C,D).**
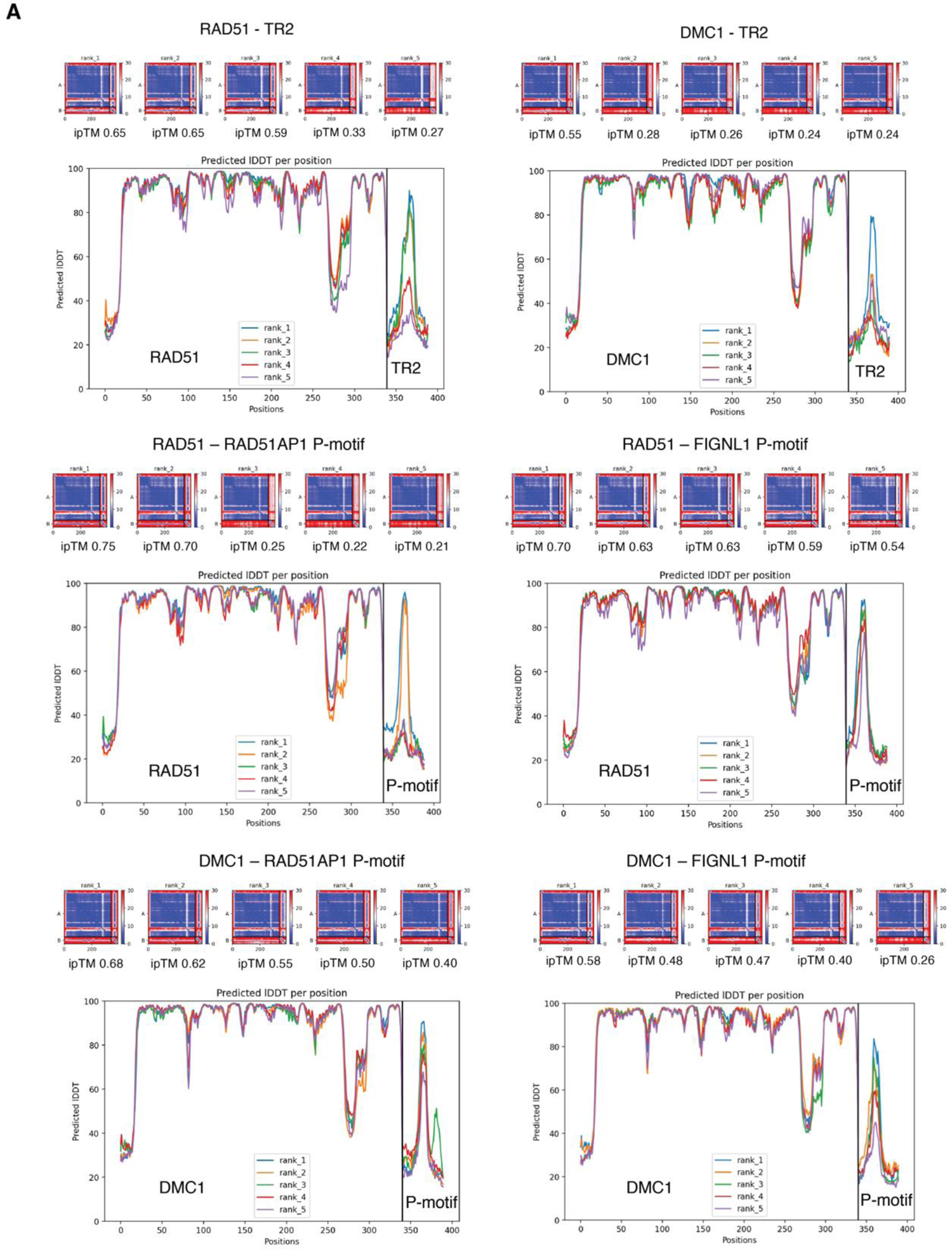
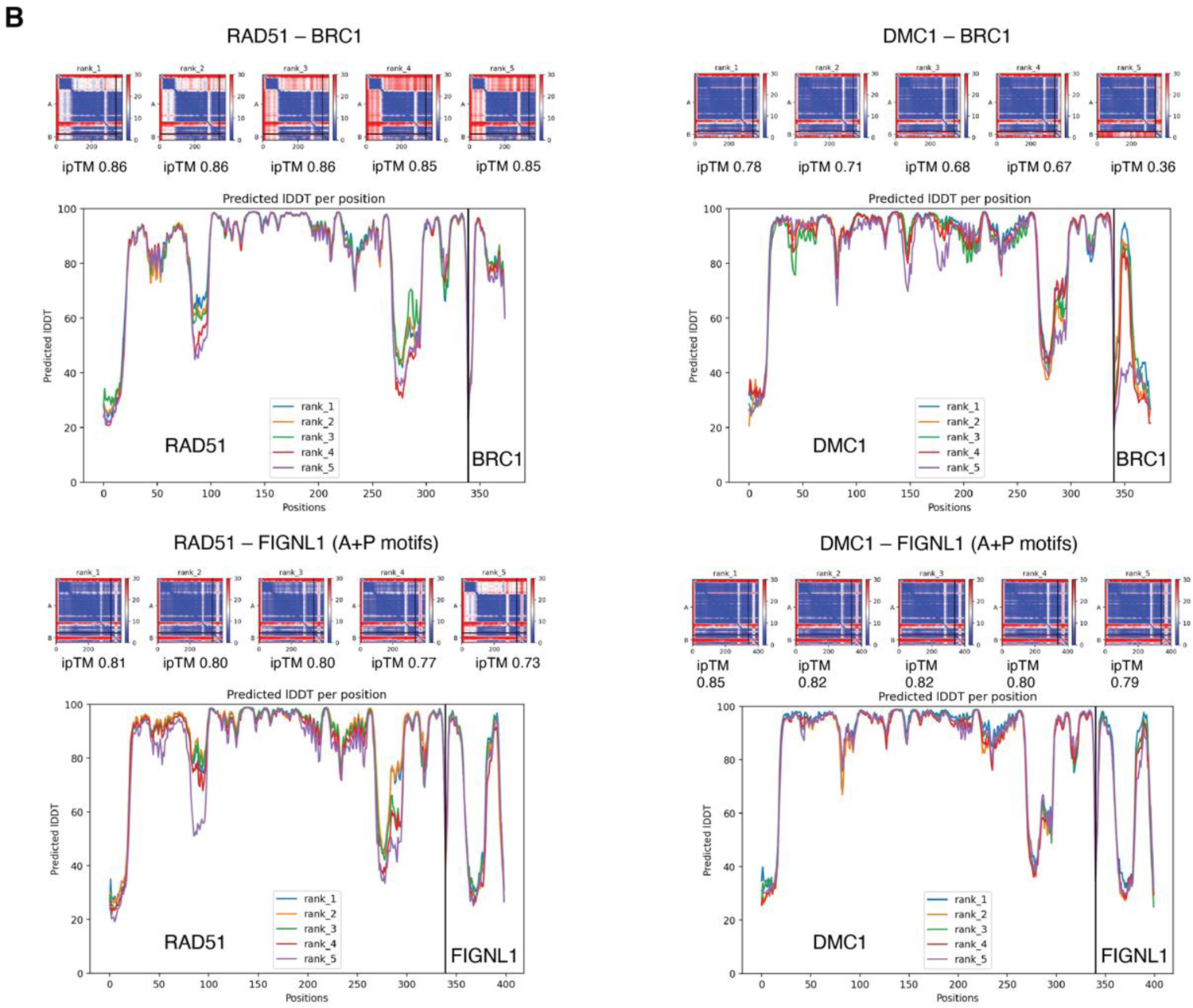
AlphaFold plots scoring the 5 models calculated for each complex: (A) scoring of the complexes from Figure 4C; (B) scoring of the complexes from Fig. 4D. In the upper panels, the heat maps show, for each of the 5 calculated models, the predicted relative position error (in Å) calculated between all pairs of residues (DMC1 and P-motif residues are numbered as 1-340 and 341-390, respectively; the color scale goes from blue - no error - to red - error above 30 Å). In the lower panels, the lDDT score is plotted as a function of the residue number.

**Supplementary Figure S5 (related to Figure 4F,G).**
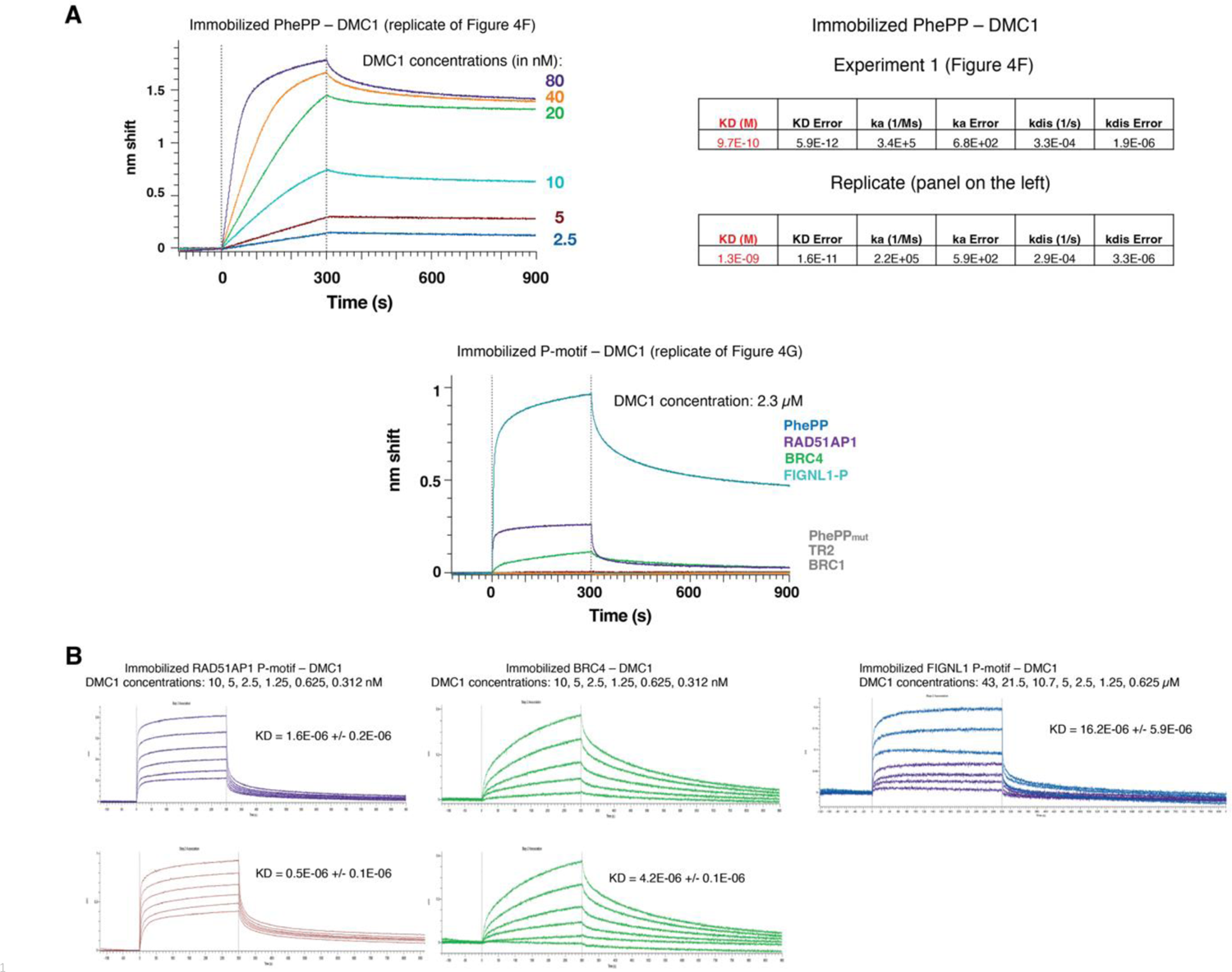
BioLayer Interferometry curves corresponding to (A) replicates of Figures 4F, G and (B) measurement of the apparent affinities between DMC1 and each motif.

**Supplementary Figure S6.**
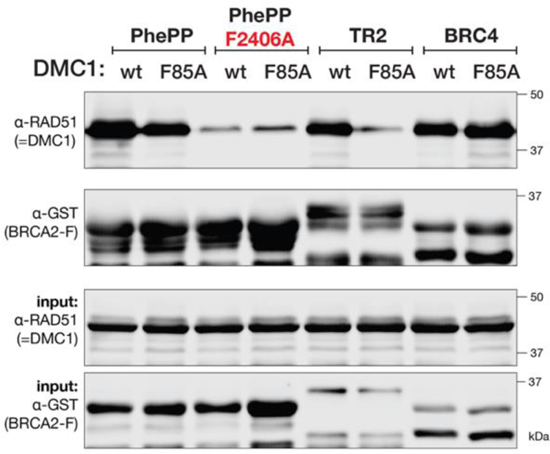
Replicates of the experiments shown in Figure 5B.

**Supplementary Figure S7.**
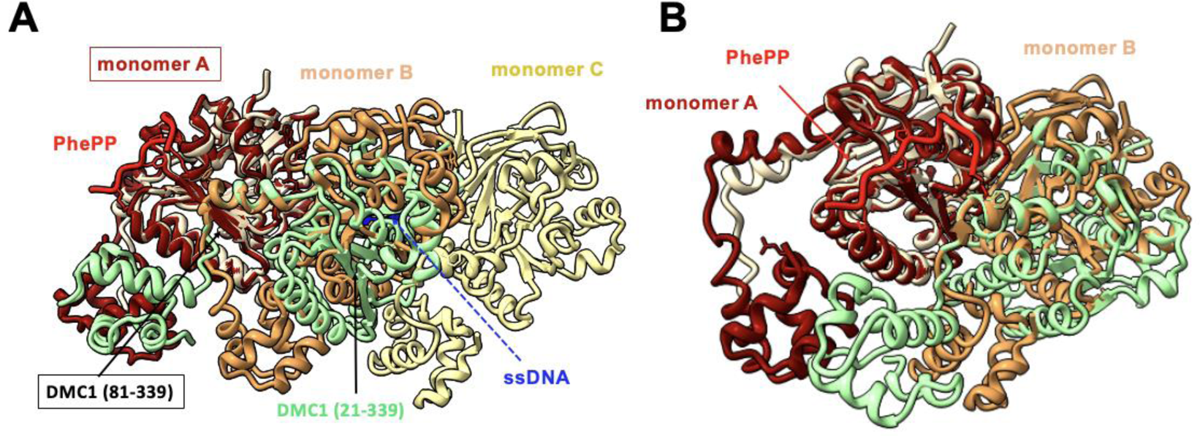
(A) Superimposition of the crystal structure of DMC1-PhePP, represented as in Figure 5A, onto the cryo-EM structure of 3 DMC1 monomers (in red, orange and yellow) assembled onto a ssDNA (in blue; PDB 7C9C). The crystal structure of DMC1 (81-339; in wheat) was fitted onto the cryo-EM structure of monomer A (in red). Consequently, the structures of DMC1 (28-339; in green) and monomer B (in orange) are shifted. This is due to the slightly different relative orientations of DMC1 monomers in the octamer and nucleoprotein filament. Despite this difference, the local environment of the PhePP residues is similar in the crystal structure and in the DMC1-PhePP-ssDNA model. (B) Zoom from (A), showing that the PhePP peptide might similarly interact with the DMC1 ATPase domain when bound to both DMC1 oligomers, but is positioned differently relatively to the DMC1 N-terminal domain when docked onto the DMC1-ssDNA filament. The N-terminal domain of the green DMC1 monomer that was close to the peptide bound to the wheat monomer is shifted in the filament model, but the N-terminal of monomer A is now close to the peptide and might similarly indirectly contribute to the peptide binding. In this panel, monomer C and the ssDNA have been hidden for clarity.

---

**Table S1.** Crystallographic data and refinement parameters for the crystal structure of DMC1-PhePP.

|  |  |
| --- | --- |
| PDB code | **8QQE** |
| Space group | *I*432 |
| Cell parameters (Å) | *a* = *b* = *c* = 234.38 |
| Resolution (Å) | 50. - 3.46<br>(3.55 - 3.46) |
| No. of observed reflections | 1119432 (80774) |
| No. of unique reflections | 14667 (1058) |
| Completeness (%) | 99.9 (99.9) |
| *R*<sub>merge</sub> (%) | 22.8 (634.5) |
| *R*<sub>pim</sub> (%) | 2.6 (72.8) |
| *I*/σ(*I*) | 22.9 (1.1) |
| *CC*<sub>1/2</sub> | 1.000 (0.541) |
| *R*<sub>cryst</sub> (%) | 24.85 |
| *R*<sub>free</sub> (%) | 26.09 |
| rms bond deviation (Å) | 0.006 |
| rms angle deviation (°) | 0.80 |
| Average *B* (Å<sup>2</sup>) |  |
| All protein atoms | 169.0 |
| By chain: |  |
| DMC1 -A-Nterm (28-84) | 247.5 |
| DMC1 -A-Cterm (85-339) | 165.9 |
| DMC1 -B-Cterm (85-339) | 153.9 |
| Peptide PhePP -C | 161.2 |
| Peptide PhePP -D | 220.0 |
| <sup>a</sup>Clashscore | 5.58 |
| MolProbity score | 1.87 |
| <sup>a</sup>Ramachandran plot (%) |  |
| Favored | 97.6 |
| Outliers | 0.2 |

Values for the highest resolution shell are in parentheses. *CC*<sub>1/2</sub>= percentage of correlation between intensities from random half-dataset. <sup>a</sup>Calculated with MolProbity

